# Zα2 domain of ZBP1 is a molecular switch regulating Influenza-induced PANoptosis and perinatal lethality during development

**DOI:** 10.1101/2020.04.05.026542

**Authors:** Sannula Kesavardhana, R. K. Subbarao Malireddi, Amanda R. Burton, Shaina N. Porter, Peter Vogel, Shondra M. Pruett-Miller, Thirumala-Devi Kanneganti

**Affiliations:** Department of Immunology, St. Jude Children’s Research Hospital, Memphis, TN, 38105, USA; Animal Resources Center and the Veterinary Pathology Core, St. Jude Children’s Research Hospital, Memphis, TN, 38105, USA; Center for Advanced Genome Engineering (CAGE), St. Jude Children’s Research Hospital, Memphis, TN, 38105, USA

**Keywords:** ZBP1, Zα domain, Z-nucleic acid, PANoptosis, influenza A virus, inflammasome, pyroptosis, apoptosis, necroptosis, RIPK1, RHIM, NLRP3

## Abstract

Z-DNA-binding protein 1 (ZBP1) is an innate nucleic acid sensor which regulates host defense responses and development. ZBP1 activation triggers inflammation and pyroptosis, necroptosis, and apoptosis (PANoptosis) by activating RIPK3, caspase-8, and the NLRP3 inflammasome. ZBP1 is unique among innate sensors because of its N-terminal Zα1 and Zα2 domains, which bind to nucleic acids in the Z-conformation. However, the specific role of these Zα domains in orchestrating ZBP1 activation and subsequent inflammation and cell death is not clear. Here we have generated *Zbp1*^Δ*Z*α2/Δ*Z*α2^ mice that lack the Zα2 domain of ZBP1 and demonstrate that this domain is critical for influenza A virus (IAV)-induced PANoptosis and perinatal lethality in RIPK1-RHIM mutated (*Ripk1*^RHIM/RHIM^) mice. Deletion of the Zα2 domain in ZBP1 abolished IAV-induced PANoptosis and NLRP3 inflammasome activation. Furthermore, deletion of the Zα2 domain of ZBP1 was sufficient to rescue *Ripk1*^RHIM/RHIM^ mice from the perinatal lethality which is caused by ZBP1-driven cell death and inflammation. Our findings identify the essential role of the Zα2 domain of ZBP1 in physiological functions and establish a link between sensing of Z-RNAs via the Zα2 domain and the promotion of influenza-induced PANoptosis and perinatal lethality.

## INTRODUCTION

Activation of inflammation and cell death are interconnected cellular events which regulate host defense and immune responses. RIP homotypic interaction motif (RHIM)-family proteins play a major role in orchestrating inflammation and cell death to profoundly shape immune responses and host defense (1,2). Z-DNA-binding protein 1 (ZBP1) is one such RHIM protein that interacts with the RHIM proteins receptor-interacting protein kinase 3 (RIPK3) and receptor-interacting protein kinase 1 (RIPK1) by forming homotypic interactions (3). ZBP1 and RIPK3 were also shown to participate in virus-induced necroptosis (4). We recently demonstrated that ZBP1 recognizes infection with influenza A virus (IAV), an RNA virus, and triggers NLRP3 inflammasome activation and pyroptosis, necroptosis, and apoptosis (PANoptosis) pathways (5). ZBP1 activation assembles RIPK3 and caspase-8 signaling complexes which mediate activation of the NLRP3 inflammasome and pyroptosis, RIPK3–MLKL-driven necroptosis, and FADD–caspase-8-driven apoptosis (5–8). These studies unraveled specific cellular functions that are regulated by ZBP1 in response to influenza infection. Further studies identified an endogenous physiological function of ZBP1 by demonstrating its role in perinatal lethality in mice (9,10). These studies found that mice with mutations that abolish the RHIM activity of RIPK1 have perinatal lethality and skin inflammation, and activation of ZBP1-driven necroptosis is responsible for the pathological phenotypes seen in these mice (9,10).

ZBP1 consists of two N-terminal Z-nucleic acid-binding domains (Zα1 and Zα2) followed by RHIM domains and a functionally undefined C-terminal region (11). The Zα1 and Zα2 domains of ZBP1 bind to DNA/RNA in the Z-conformation (12–15). Recognition of endogenous RNA, viral RNAs, and vRNPs that might exist in the Z-nucleic acid conformation is thought to activate ZBP1-mediated inflammation and cell death (6,8,16). The Zα domains of ZBP1 show high affinity for Z-nucleic acids *in vitro* (12–14,17); however, whether the recognition of Z-nucleic acids regulates cellular functions is not studied. In addition, the role of the Z-nucleic acid-binding Zα domains of ZBP1 in regulating cell death and inflammation remains poorly investigated. Here we identify the critical role of the Zα2 domain of ZBP1 in regulating IAV-induced NLRP3 inflammasome activation, cell death, and perinatal lethality during mouse development. Our observations define a pivotal role for the Zα2 domain of ZBP1 *in vivo* and provide evidence for the activation of ZBP1 by endogenous Z-RNAs.

## RESULTS

### Generation of C57BL/6 mice that lack the Zα2 domain of ZBP1

Experiments using ZBP1-knockout mice, which show complete loss of ZBP1 expression throughout the body (*Zbp1^−/−^*), have delineated the role of ZBP1 during *in vivo* infections and in mouse development (11). ZBP1 is a critical innate immune sensor for activating inflammatory signaling and PANoptosis in both immune and non-immune cells in response to IAV infection (5,18). In addition, ZBP1 also activates necroptosis and inflammation during RIPK1-RHIM mutant mouse development, leading to perinatal lethality (6,9,10). Both the Zα1 and Zα2 domains of ZBP1 are capable of binding to Z-nucleic acids, and specific mutations in either of these Zα domains block their Z-nucleic acid-binding activity (12,13,19). Although several recent studies established the role of ZBP1 in innate immunity and development, specific contributions from the Z-nucleic acid-recognizing Zα domains of ZBP1 *in vivo* are not clear. To investigate this, we generated ZBP1 mutant mice with deletion of the Zα2 domain in the C57BL/6 background using CRISPR/Cas9 technology (hereafter referred as *Zbp1*^ΔZα2/ΔZα2^ mice) (**Fig. 1A** **and** **Suppl Table 1**). At the amino acid sequence level, the CRISPR/Cas9 approach generated a Zα2 domain deletion in ZBP1 by joining residues at positions 76 and 151 and deleting residues at positions 77–150 (**Fig. 1A**). *Zbp1*^ΔZα2/ΔZα2^ mice were viable and did not show visible phenotypic defects.

**Figure 1:**
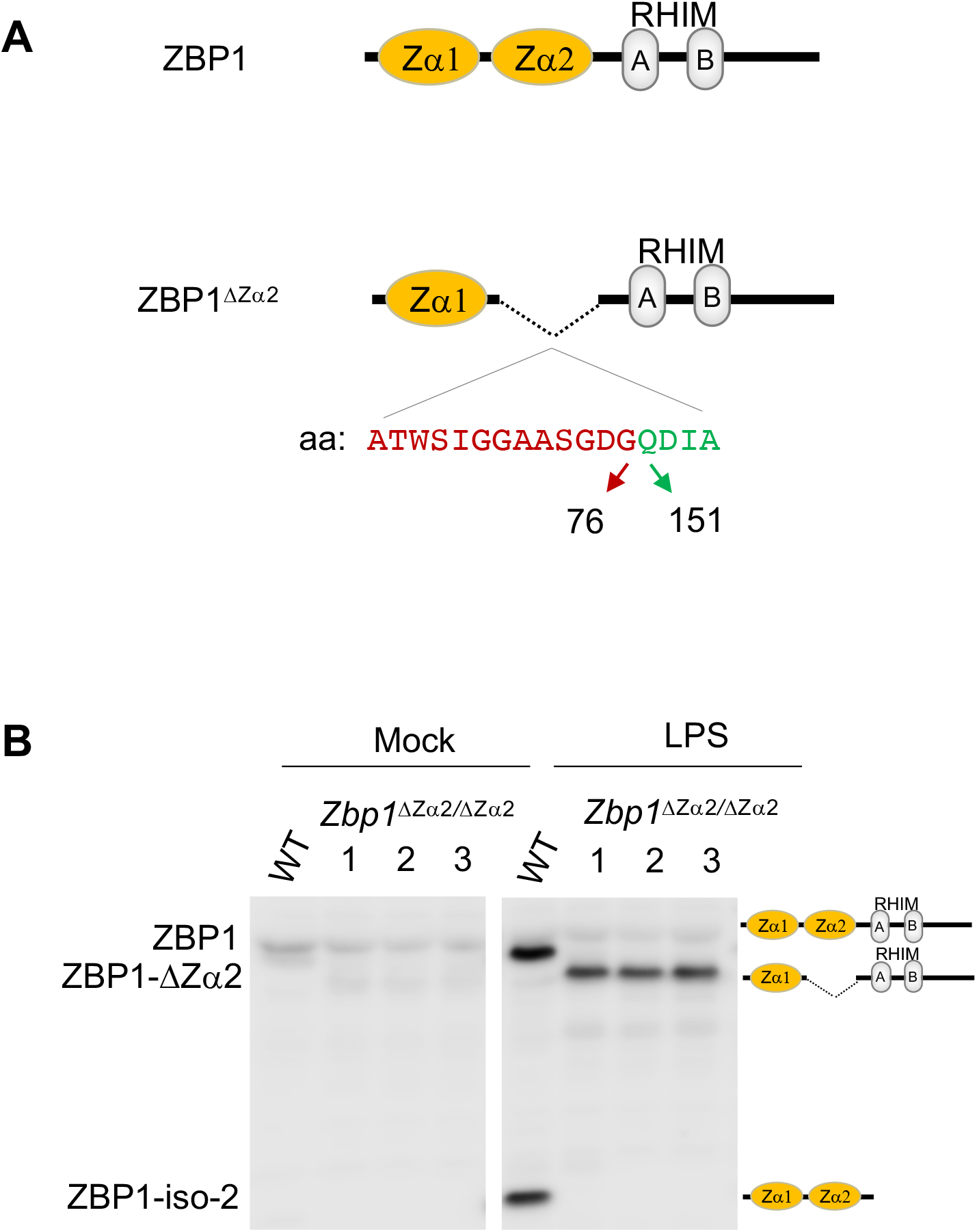
Generation of mice with Za2 domain deletion in ZBP1 (*Zbp1* ^ΔZα2/ΔZα2^) **A,** Schematic representation of domain structure and deletion of the Zα2 domain in ZBP1. Amino acid residues present at the N-terminus (red) and C-terminus (green) of the Zα2 domain which are in sequence after the deletion of this domain are represented. aa, amino acid residues. **B,** Immunoblot analysis of ZBP1 after mock or LPS stimulation of WT and *Zbp1* ^ΔZα2/ΔZα2^ BMDMs. ZBP1-ΔZα2, ZBP1 with deletion of the Zα2 domain; ZBP1-iso-2, ZBP1 isoform 2.

To confirm the deletion of the Zα2 domain at the protein level, we generated bone marrow-derived macrophages (BMDMs). *Zbp1*^ΔZα2/ΔZα2^ BMDMs showed similar expression of the myeloid markers CD11b and F4/80 compared with BMDMs generated from WT littermates, suggesting the deletion caused no defects in the ability of bone marrow progenitors to differentiate into macrophages (**Suppl Fig. 1**). Lipopolysaccharide (LPS) stimulation induced upregulation of ZBP1 expression in both WT and *Zbp1*^ΔZα2/ΔZα2^ BMDMs, indicating the Zα2 domain deletion did not interfere with the expression of ZBP1 protein (**Fig. 1B**). A shift in ZBP1 protein towards a lower molecular weight was observed in *Zbp1*^ΔZα2/ΔZα2^ BMDMs, confirming the deletion of the Zα2 domain (ZBP1-ΔZα2) in these mice (**Fig. 1B**). Isoform-2 of ZBP1, which consists of only the Zα1-Zα2 domains, was not seen in *Zbp1*^ΔZα2/ΔZα2^ BMDMs (**Fig. 1B**). These results established the deletion of the Zα2 domain of ZBP1 in mice with no detectable phenotypic differences in the mice and macrophages.

### Zα2 domain of ZBP1 triggers inflammasome activation and PANoptosis during IAV infection

ZBP1 plays a crucial role in activation of the NLRP3 inflammasome and PANoptosis upon recognizing IAV infection (5,6). To understand the role of the Zα2 domain in ZBP1-induced NLRP3 inflammasome activation and PANoptosis, we infected WT and *Zbp1*^ΔZα2/ΔZα2^ BMDMs with mouse-adapted IAV (influenza A/Puerto Rico/8/34 [PR8; H1N1]). Similar to LPS stimulation, IAV infection upregulated the expression of ZBP1-ΔZα2 to a similar extent that ZBP1 was upregulated (**Fig. 2A**). In addition, there was no defect in IAV replication in *Zbp1*^ΔZα2/ΔZα2^ BMDMs as indicated by NS1 protein expression (**Fig. 2A**). IAV infection triggers specific activation of the NLRP3 inflammasome (5,6,20,21). *Zbp1*^ΔZα2/ΔZα2^ BMDMs showed a complete lack of caspase-1 (CASP1) activation compared with WT BMDMs, suggesting inhibition of IAV-induced NLRP3 inflammasome activation in these BMDMs (**Fig. 2A**). Cleavage of gasdermin D (GSDMD) and release of the GSDMD N-terminal domain (p30), which is a measure of GSDMD activation, was also inhibited in *Zbp1*^ΔZα2/ΔZα2^ BMDMs in response to IAV infection (**Fig. 2A**). These observations suggest that the Zα2 domain is critical for ZBP1-mediated activation of NLRP3 inflammasome and GSDMD-induced pyroptosis.

**Figure 2:**
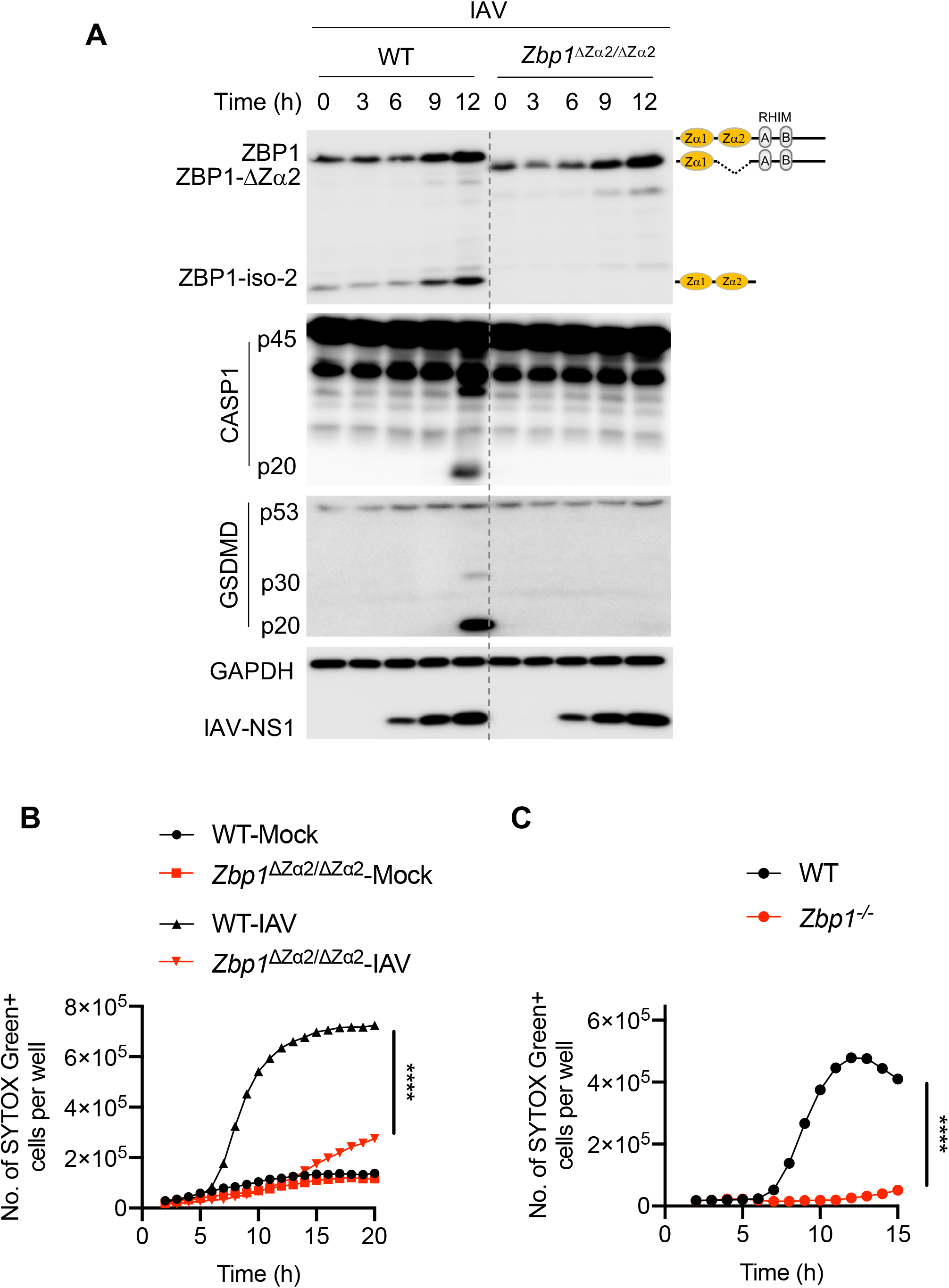
Zα2 domain of ZBP1 is critical for triggering IAV-induced NLRP3 inflammasome activation and PANoptosis. **A,** Immunoblot analysis of ZBP1, caspase-1 (CASP1), gasdermin D (GSDMD), IAV-NS1, and GAPDH in WT and *Zbp1* ^ΔZα2/ΔZα2^ BMDMs infected with IAV (mouse adapted, influenza A/Puerto Rico/8/34 [PR8; H1N1]). **B and C,** Cell death as measured by the number of SYTOX Green^+^ cells. BMDMs were infected with IAV, and cell death was monitored at regular intervals. \*\*\*\**P* < 0.0001 (two-way ANOVA).

We further investigated the role of the Zα2 domain of ZBP1 in regulating IAV-induced PANoptosis in BMDMs. We found that IAV infection induced robust cell death in WT BMDMs compared with mock treatment (**Fig. 2B**). This IAV-induced cell death was completely abolished in *Zbp1*^ΔZα2/ΔZα2^ BMDMs, suggesting a lack of ZBP1-mediated signaling (**Fig. 2B**). Consistent with previous reports (5,6,8), *Zbp1*^−/−^ BMDMs showed complete inhibition of IAV-induced cell death (**Fig. 2C**). These results suggest that the Zα2 domain of ZBP1 is necessary and sufficient for ZBP1 sensing of IAV infection to engage PANoptosis. Deletion of the Zα2 domain of ZBP1 showed defects in cellular functions similar to those seen with the loss of complete ZBP1 expression.

### Zα2 domain is crucial for ZBP1-induced necroptosis and perinatal lethality in RIPK1-RHIM mutant mice

Previous studies unraveled an important role for ZBP1 by studying RIPK1-RHIM mutant mice (9,10). Mutations in the RHIM domain of RIPK1 (*Ripk1*^RHIM/RHIM^) cause perinatal lethality in mice. This perinatal lethality is driven by ZBP1-induced activation of RIPK3– MLKL-mediated necroptosis and inflammation. Deletion of ZBP1 or RIPK3 rescues *Ripk1*^RHIM/RHIM^ mice from excessive cell death and perinatal lethality. These findings demonstrate the role of ZBP1 in regulating cell death under physiological conditions and during mouse development. However, how ZBP1 is activated in *Ripk1*^RHIM/RHIM^ mice and whether sensing of endogenous Z-RNAs via Zα domains triggers perinatal lethality is not clear. To understand this, we tested whether deletion of the Zα2 domain of ZBP1 is sufficient to rescue *Ripk1*^RHIM/RHIM^ mice from perinatal lethality. Consistent with recent studies (9,10), deletion of ZBP1 rescued *Ripk1*^RHIM/RHIM^ mice from lethality, and viable mice were only produced when both the alleles of *Zbp1* were absent (**Suppl Fig. 2A**). We found that deletion of the Zα2 domain of ZBP1 also rescued perinatal lethality in *Ripk1*^RHIM/RHIM^ mice, and the mice became viable (**Fig. 3A** **&** **3B**). Similar to WT and *Ripk1*^RHIM/RHIM^ *Zbp1*^−/−^ mice, *Ripk1*^RHIM/RHIM^ *Zbp1* ^ΔZα2/ΔZα2^ mice also survived and developed normally into adulthood (**Fig. 3C**). Deletion of the Zα2 domain in both the alleles of *Zbp1* was necessary to make *Ripk1*^RHIM/RHIM^ mice viable (**Fig. 3A–3C**). This suggests that complete inhibition of the Z-RNA sensing activity of ZBP1 through deletion of the Zα2 domain is necessary to inhibit ZBP1-driven necroptosis and perinatal lethality in *Ripk1*^RHIM/RHIM^ mice. These observations identify that the Zα2 domain of ZBP1 is critical for driving cell death during development in *Ripk1*^RHIM/RHIM^ mice which in turn indicates potential sensing of endogenous RNA ligands by ZBP1.

**Figure 3:**
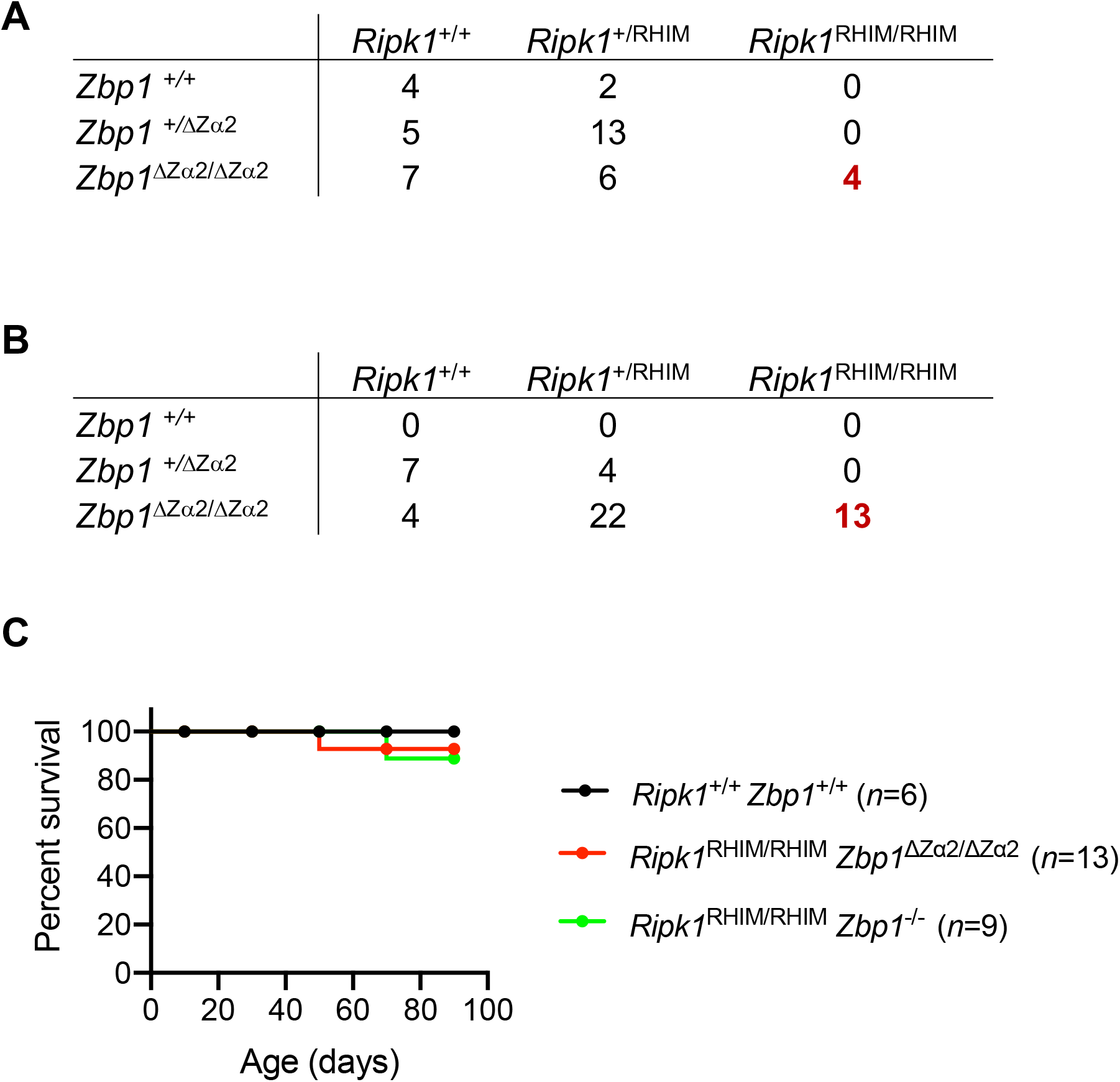
Z-nucleic acid-binding Zα2 domain of ZBP1 is essential for embryonic development and acts as an activator of RIPK1-RHIM mutation-induced perinatal lethality in mice. **A,** Table representing the number of offspring of each genotype generated from intercrossing *Ripk1*^+/RHIM^ *Zbp1* ^+/ΔZα2^ (heterozygote) parents. **B,** Table representing the number of offspring of each genotype generated from intercrossing *Ripk1*^+/RHIM^ *Zbp1* ^ΔZα2/ΔZα2^ parents. In both **A** and **B,** the numbers of *Ripk1*^RHIM/RHIM^ mice rescued from perinatal lethality are highlighted in red. **C,** Kaplan-Meier plot showing survival of *Ripk1*^RHIM/RHIM^ *Zbp1* ^ΔZα2/ΔZα2^ mice and *Ripk1*^RHIM/RHIM^ *Zbp1*^−/−^ mice compared with WT *Ripk1*^+/+^ *Zbp1* ^+/+^ mice.

To further investigate the impact of deleting the Zα2 domain of ZBP1, we studied the immune cell population in circulation. The numbers and percentages of circulating lymphocytes, monocytes, and red blood cells were unaltered in viable *Ripk1*^RHIM/RHIM^ *Zbp1* ^ΔZα2/ΔZα2^ mice in comparison to WT mice (**Suppl Fig. 2B**). We observed a higher percentage of circulating neutrophils in both *Ripk1*^RHIM/RHIM^ *Zbp1* ^ΔZα2/ΔZα2^ mice and *Ripk1*^RHIM/RHIM^ *Zbp1*^−/−^ mice (**Suppl Fig. 2B**). Consistent with these observations, a previous study reported neutrophilic inflammation in *Ripk1*^RHIM/RHIM^ *Zbp1*^−/−^ mice (10). Ex vivo differentiated BMDMs from *Ripk1*^RHIM/RHIM^ *Zbp1* ^ΔZα2/ΔZα2^ mice and *Ripk1*^RHIM/RHIM^ *Zbp1*^−/−^ mice showed similar CD11b and F/480 myeloid expression markers in comparison to WT BMDMs (**Suppl Fig. 2C**). In addition, IAV infection did not induce cell death in both *Ripk1*^RHIM/RHIM^ *Zbp1* ^ΔZα2/ΔZα2^ and *Ripk1*^RHIM/RHIM^ *Zbp1*^−/−^ BMDMs (**Suppl Fig. 2D**). These results suggest that the Zα2 domain of ZBP1 prevented cell death and perinatal lethality in *Ripk1*^RHIM/RHIM^ mice and facilitated normal immune cell development in these mice, with a mild increase in circulating neutrophils.

## DISCUSSION

Several recent studies unraveled an important role for ZBP1 in regulating cell death and inflammation and uncovered its role as an innate immune sensor. ZBP1 was first described as a tumor-associated protein that was upregulated in macrophages (22). Identification of a RHIM domain in ZBP1 facilitated its characterization in cell death and inflammation (3,4,11). The RHIM-containing proteins regulate inflammatory signaling, apoptosis, and inflammatory cell death (1). Initial reports on ZBP1 suggested it plays a role in virus-induced necroptosis via its RHIM-mediated interaction with RIPK3 (3,4). We identified that IAV infection and its RNA genome products activate the NLRP3 inflammasome to mount inflammation and lung repair responses *in vivo* (20,21,23). Identification of ZBP1’s role in sensing IAV and activating the inflammasome and multiple programmed cell death pathways expanded its functions in diverse cellular activities and illustrated potential regulation of inflammation *in vivo* (5,11). The N-terminal Zα domains of ZBP1 bind to nucleic acids in the Z-conformation. The Zα domains bind to both Z-DNA or Z-RNA because of their conformational similarity once they attain the Z-conformation (12,13,15,17,19). Although Zα domains interact with Z-nucleic acids *in vitro*, activation and cellular functions of these Zα domains of ZBP1 and *in vivo* relevance were not studied.

Our findings here suggest that the Zα2 domain is critical for the activation of ZBP1’s functions. The Zα2 domain is required for activation of ZBP1 to drive NLRP3 inflammasome activation, PANoptosis, and perinatal lethality in mice. It is interesting to find that deletion of the Zα2 domain is sufficient to block ZBP1’s functions. Our observations also suggest that the Zα1 domain might be dispensable for ZBP1 regulation *in vivo*. The Zα2 domain of ZBP1 shows a distinct way of binding to Z-nucleic acids unlike other Zα domains from various other proteins (12). This suggests that the Zα2 domain of ZBP1 might have evolved to control cellular functions by executing distinct mechanisms, and our findings showing its critical role in regulating host innate responses support this hypothesis. Earlier *in vitro* studies also demonstrated that deletion of the Zα1 domain was not sufficient to inhibit ZBP1-induced cell death (8). Our results also suggest that the Zα2 domain of ZBP1 may recognize viral or endogenous RNAs which attain Z-conformation to activate ZBP1-driven cell death and perinatal lethality. The N122 and Y126 residues in the Zα2 domain of ZBP1 are essential for the Z-nucleic acid interaction, and mutations in these positions were shown to abolish the Z-nucleic acid-binding potential of ZBP1 (12). Thus, the Zα2 domain of ZBP1 is a crucial determinant of ZBP1 for sensing Z-nucleic acids and subsequent engagement of inflammation and cell death. Observations from IAV infection experiments suggest that deletion of the Zα2 domain phenocopied complete ZBP1 deletion in regulating PANoptosis and the activation of the NLRP3 inflammasome. While this manuscript was in preparation, new studies appeared demonstrating that sensing of influenza or endogenous retroviral Z-nucleic acids via the Zα domains of ZBP1 is essential for activating cell death and inflammation (24–26). These findings from independent groups corroborate our results presented here and also our previous reports on ZBP1 sensing of an RNA virus (IAV) to activate cell death and inflammation (5,6,24–26). Overall, our findings, in conjunction with these new studies, establish a critical and specific role for the Zα2 domain of ZBP1 in regulating PANoptosis, inflammation, and perinatal lethality in mice and provide genetic evidence for the functional relevance of Z-nucleic acid-binding domains *in vivo*.

## EXPERIMENTAL PROCEDURES

### Mice

The Zα2 domain of ZBP1 was deleted in C57BL/6 mice (*Zbp1* ^ΔZα2/ΔZα2^) using CRISPR/Cas9 gene editing. Briefly, C57BL/6J (Jackson Laboratories, Bar Harbor, ME) fertilized zygotes were injected into the pronucleus with 15 ng/μL of each chemically modified sgRNA (Synthego), 60 ng/μL SpCas9 protein (Berkeley Microlab), and 10 ng/μL ssODN donor (IDT) diluted in 0.1 mM EDTA. Founder mice were genotyped by targeted next generation sequencing on a MiSeq Illumina Sequencer and analyzed with CRIS.py (27). Editing construct sequences and relevant primers are listed in **Suppl. Table 1**. After confirming the deletion, mice were backcrossed to C57BL/6 genetic background mice at least three times. *Ripk1*^+/RHIM^ mice were a gift from Dr. Vishva M Dixit from Genentech (10). These mice were crossed to *Zbp1* ^ΔZα2/ΔZα2^ mice and *Zbp1*^−/−^ mice to generate *Ripk1*^RHIM/RHIM^ *Zbp1* ^ΔZα2/ΔZα2^ mice and *Ripk1*^RHIM/RHIM^ *Zbp1*^−/−^ mice. Offspring were monitored for survival and rescue of perinatal lethality in *Ripk1*^RHIM/RHIM^ mice. Mutations or deletions were confirmed by PCR. All mice were bred and housed in the Animal Resource Center at St. Jude Children’s Research Hospital. Animal study protocols were approved by St. Jude Children’s Research Hospital committee on the care and use of animals.

### BMDM cultures and IAV infection

Primary BMDMs from mouse bone marrow were grown for 6 days in IMDM (Gibco) media supplemented with 30% L929-conditioned media, 10% FBS (Atlanta Biologicals), 1% penicillin and streptomycin, and 1% non-essential amino acids (Gibco). BMDMs were seeded and cultured overnight before using them for stimulations or infections. For lipopolysaccharide (LPS) stimulation experiments, BMDMs were mock treated (media only) or stimulated with 100 ng/mL of LPS (InvivoGen) for 4 h. The IAV-PR8 strain was generated as described previously (5,6). BMDMs were infected with IAV (MOI, 10) for 2 h in serum-free DMEM media (Sigma). After 2 h of infection, 10% FBS was added. Whole cell lysates were collected at indicated time points for immunoblotting analysis or cell death was monitored.

## Immunoblotting analysis

Immunoblotting and detection was performed as described previously (6). BMDMs were washed with PBS once, cells were lysed in RIPA buffer, and sample loading buffer containing SDS and 2-mercaptoethanol was added. Proteins were separated on 8%–12% polyacrylamide gels and transferred onto PVDF membranes (Millipore). Membranes were blocked in 5% skim milk followed by incubation with primary antibodies and secondary horseradish peroxidase (HRP)-conjugated antibodies. Primary antibodies used were anti-ZBP1 (AG-20B-0010-C100, AdipoGen), anti–caspase-1 (AG-20B-0042, AdipoGen), anti-GSDMD (Ab209845, Abcam), anti-GAPDH (5174, CST), and anti-IAV NS1 (sc-130568, Santa Cruz Biotechnology Inc.). HRP-conjugated secondary antibodies (anti-rabbit [111-035-047], anti-mouse [315-035-047], Jackson Immuno Research Laboratories) were used. Protein detection was done by using Luminata Forte Western HRP Substrate (Millipore, WBLUF0500).

### SYTOX Green staining and cell death analysis

Real-time cell death assays of IAV-infected BMDMs were performed using an IncuCyte Zoom incubator imaging system (Essen Biosciences). BMDMs were seeded in 12-well tissue culture plates in the presence of 100 nM SYTOX Green (Thermo Fisher Scientific, S7020), which is a cell-impermeable DNA-binding fluorescent dye that enters dying cells upon membrane permeabilization. Analysis was done using the software package supplied with the IncuCyte imager. Using this software, a precise analysis of the number of SYTOX Green-positive (SYTOX Green+) cells present in each image can be performed. The number of dead cells was acquired by counting SYTOX Green+ cells and plotted using GraphPad Prism v8 software.

### Flow cytometry

Flow cytometry analysis was performed to monitor differentiation of bone marrow cultures into macrophages. For analysis, the following monoclonal antibodies were used: anti-mouse CD45.2 (Clone 104; BioLegend, 109806), anti-CD11b (M1/70; Affymetrix eBioscience, 48-0112-82), and anti-F4/80 (BM8; BioLegend, 123116). These antibodies were diluted 1:300 to stain BMDMs for 15 min at 4°C in FACS buffer (2% v/v of FBS in phosphate-buffered saline containing 0.04% w/v sodium azide). Flow cytometry data were acquired on an LSR II flow cytometer (BD) and analyzed with FlowJo software (FlowJo, LLC, and Illumina, Inc.).

## ACKNOWLEDGEMENTS

We sincerely thank Dr. Vishva M Dixit from Genentech for sharing *Ripk1*^+/ RHIM^ mice. We also thank Rebecca Tweedell, PhD, for scientific editing. This study was supported by grants from the US National Institutes of Health (AR056296, CA163507, AI101935, and AI124346) and the American Lebanese Syrian Associated Charities to T.D.-K.

## AUTHOR CONTRIBUTIONS

S.K. and T.-D.K. conceptualized the study; S.K., R.K.S.M., and T.-D.K. designed the methodology; S.K., R.K.S.M., and A.R.B. performed the experiments; S.N.P. and S.M.P-M. generated mice deficient in the Zα2 domain using CRISPR/Cas9 approach; P.V. assessed the pathology in mice; S.K., R.K.S.M., and T.-D.K. conducted the analysis; S.K. and T.-D.K. wrote the manuscript with input from all authors. T.-D.K. acquired the funding and provided overall supervision.

---

**Supplementary Figure 1:**
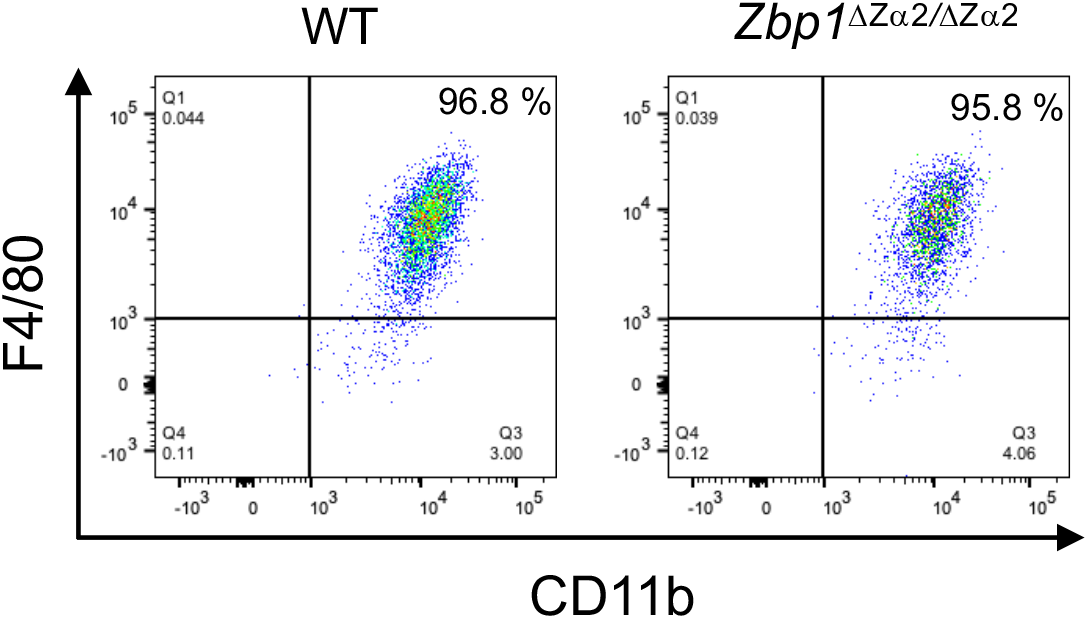
Analysis of macrophage differentiation after deletion of Zα2 domain of ZBP1. Flow cytometry analysis of ex vivo BMDMs from WT and *Zbp1* ^ΔZα2/ΔZα2^ mice for expression of myeloid-specific markers, F4/80 and CD11b.

**Supplementary Figure 2:**
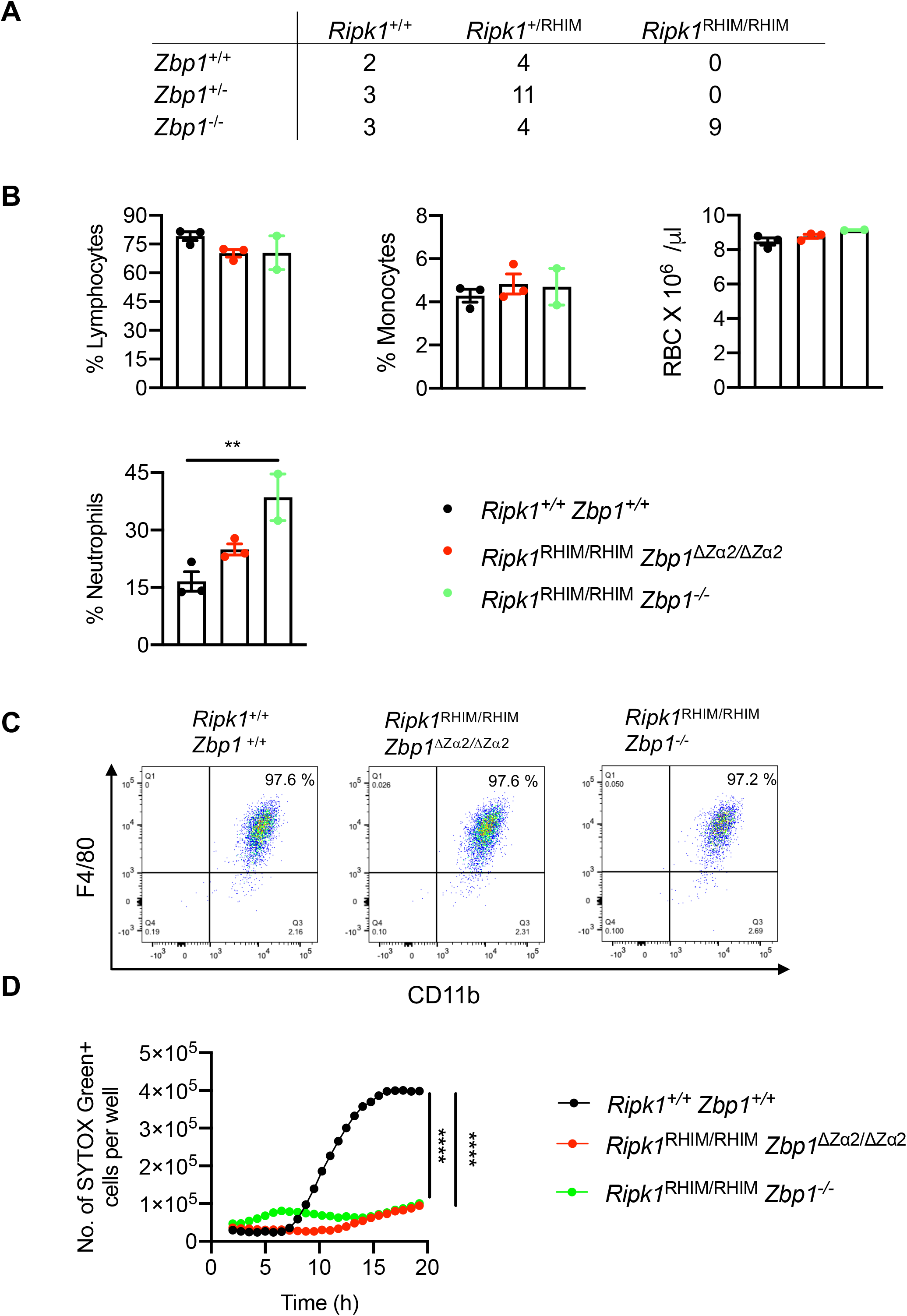
Immune cell population in *Ripk1*^RHIM/RHIM^ *Zbp1* ^Δα2/ΔZα2^ and *Ripk1*^RHIM/RHIM^ *Zbp1*^−/−^ mice and IAV-induced cell death. **A,** Table representing the number of offspring of each genotype generated from intercrossing *Ripk1*^+/RHIM^ *Zbp1*^+/−^ (heterozygote) parents. **B,** Analysis of the number or percent of different immune cell types from the blood from indicated mouse strains. Data shown are mean ± SEM. \*\**P* = 0.0462 (unpaired two-tailed *t*-test) for *Ripk1*^+/+^ *Zbp1*^+/+^ vs *Ripk1*^RHIM/RHIM^ *Zbp1* ^ΔZα2/ΔZα2^ cells; \*\**P =* 0.0296 (unpaired two-tailed *t*-test) for *Ripk1*+/+ *Zbp1*+/+ vs *Ripk1*^RHIM/RHIM^ *Zbp1*^−/−^ cells. **C,** Flow cytometry analysis of ex vivo BMDMs from the indicated mouse strains for expression of myeloid-specific markers, F4/80 and CD11b. **D,** Cell death as measured by the number of SYTOX Green^+^ cells. BMDMs differentiated from the indicated mouse strains were infected with IAV, and cell death was monitored at regular intervals. \*\*\*\**P* < 0.0001 (two-way ANOVA).

**Supplementary Table 1:**
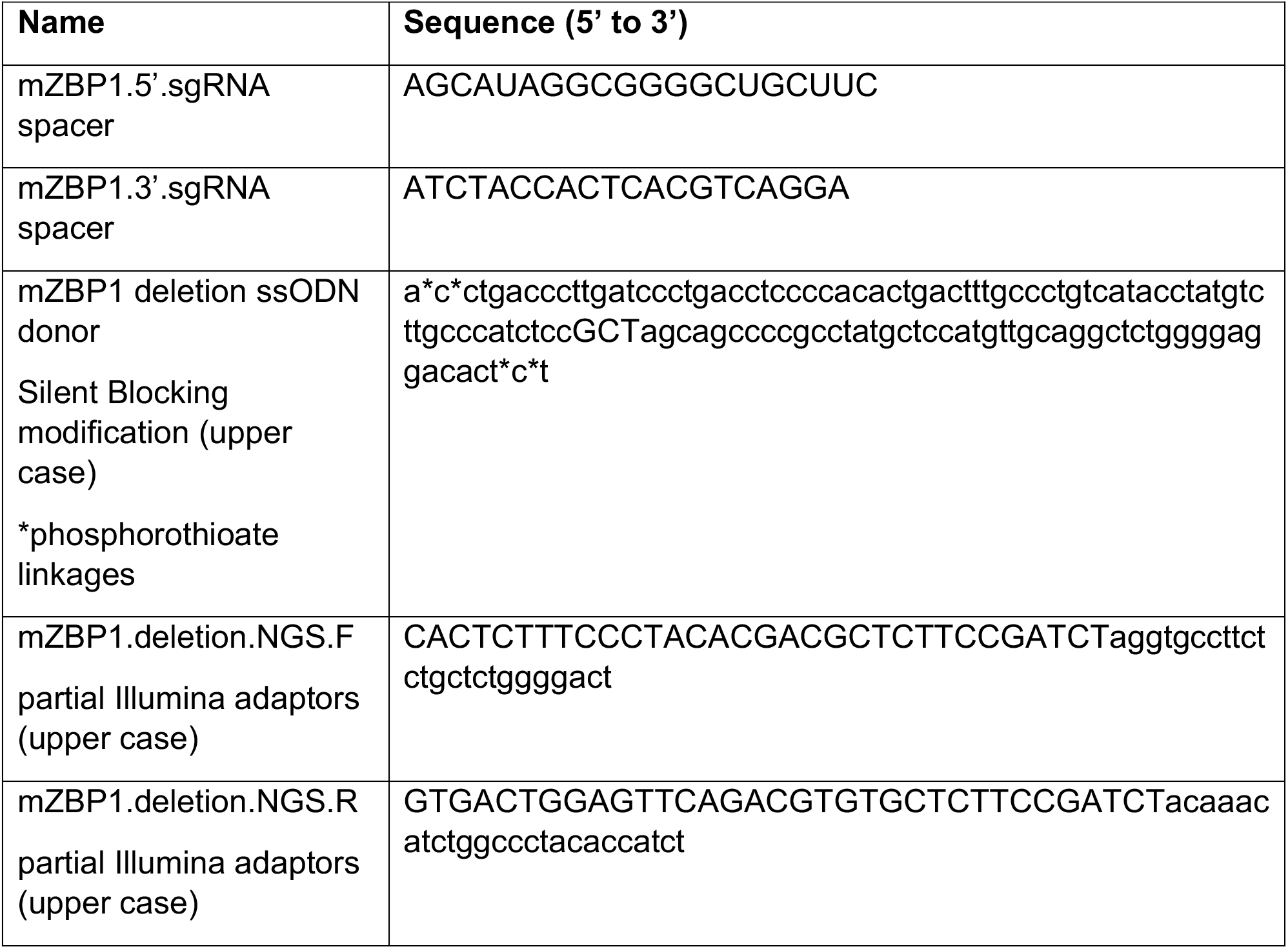
List of CRISPR/Cas9 gene editing construct sequences and relevant primers.

